# A receptor tyrosine kinase regulated by the transcription factor VAB-3/PAX6 pairs with a pseudokinase to trigger immune signalling upon oomycete recognition in *C. elegans*

**DOI:** 10.1101/2022.12.15.520564

**Authors:** Florence Drury, Manish Grover, Mark Hintze, Jonathan Saunders, Michael K Fasseas, Charis Constantinou, Michalis Barkoulas

**Affiliations:** Department of Life Sciences, Imperial College, London SW7 2AZ, United Kingdom.

**Keywords:** oomycete, pathogen recognition, innate immunity, *chil*, *Myzocytiopsis humicola*, *C. elegans*, PAX6, VAB-3, KIN-16 receptor tyrosine kinase, pseudokinase

## Abstract

Oomycetes were recently discovered as natural pathogens of *Caenorhabditis elegans* and pathogen recognition alone was shown to be sufficient to activate a protective transcriptional program in the host characterised by the expression of multiple *chitinase-like* (*chil*) genes. However, the molecular mechanisms underlying oomycete recognition in animals remain fully unknown. We performed here a forward genetic screen to uncover regulators of *chil* gene induction and found several independent loss-of-function alleles of *old-1* and *flor-1,* which encode receptor tyrosine kinases belonging to the *C. elegans*-specific KIN-16 family. We present evidence that OLD-1 is an active kinase mounting the immune response, and FLOR-1 a pseudokinase that is also required for the response and regulates the distribution of OLD-1 at the epidermal membrane. Interestingly, the *old-1* locus is adjacent to the *chil* genes in the nematode genome, thereby revealing a genetic cluster important for oomycete-resistance. Furthermore, we identify the VAB-3/PAX-6 transcription factor known for its role in visual system development to regulate *old-1* expression, and consequently the spatial pattern of the response to oomycete recognition. Taken together, our study reveals both conserved and species-specific factors shaping the response to oomycete recognition.

## Introduction

Immune responses to pathogenic infections are vital for the survival of all organisms. *Caenorhabditis elegans* is a small, free-living nematode that is used as a model to understand the molecular pathways employed by animal hosts to fight infection. *C. elegans* encounters a plethora of pathogens in its natural environment and sampling of wild isolates over the last years has allowed new natural host-pathogen interactions to be studied in the laboratory^1^. Immune responses to naturally occurring infections have co-evolved alongside the pathogen and have been studied in the context of bacterial^2,3^, viral^4^, fungal^5, 6^ and fungal-like microsporidia infections^7^. We previously described a new natural infection of *C. elegans* by the oomycete *Myzocytiopsis humicola*^8^. Oomycetes are a lesser-known eukaryotic class of pathogens that cause lethal infection in both plants^9^ and animals, including humans^10^. Human oomycete infection for example is caused by the oomycete *Pythium insidiosum*, an emerging tropical infection with a high mortality rate due to lack of awareness and suitable non-invasive treatments^11^. Therefore, *C. elegans* provides a useful model host for us to study oomycete infection in a whole-animal context.

*M. humicola* infects and rapidly kills *C. elegans* by attaching and penetrating through the cuticle and developing first into hyphae, and then sporangia inside the body cavity before the release of zoospores that go on to infect other animals^8^. Interestingly, we have also shown that *C. elegans* can detect *M. humicola* in the environment using chemosensory neurons^12^. This triggers cross-tissue communication from the neurons to the epidermis where *chitinase-like* (*chil*) genes are induced and offer some protection against infection through cuticle modification^8^. A similar kind of neuron-to-epidermis communication also exists for immune signalling against fungal infection in *C. elegans*, where neuronal expression of *dbl-1* regulates expression of *cnc* antimicrobial peptides in the epidermis^5^. The induction of *chil* genes can be elicited by an innocuous oomycete extract produced by washing off plates containing infected animals followed by filter sterilisation and autoclaving. By exposing animals to such extracts, an Oomycete Recognition Response (ORR) was described from the overlap of genes transcriptionally upregulated upon oomycete extract treatment and oomycete infection^12^. In fact, it has been previously shown that cell wall components from the oomycete *Saprolegnia parasitica*, and glucan extract from *P. insidiosum* activate Th2-like response^13^ and a specific Th1/Th17 response^14,15^ in Atlantic salmon and BALB/c mice respectively. Also, the proinflammatory response in Atlantic salmon was found to extend beyond gills in head kidney cells as well as the mucosal tissue^13^. However, the underlying mechanisms of oomycete recognition and response remain unknown, and therefore, identifying the regulators of ORR in *C. elegans*, can help better understand the response to animal oomycete infection.

In this study, we have started dissecting the underlying signalling pathway involved in oomycete recognition by identifying positive regulators of ORR. We show that Overexpression Longevity Determinant (OLD-1) and Full Loss of Oomycete response (FLOR-1), which are receptor tyrosine kinases from the KIN-16 family in *C. elegans,* are both required for mounting the ORR. We establish OLD-1 as an active kinase driving the response, while FLOR-1 as a pseudokinase that interacts with OLD-1 in the epidermal membrane and regulates both levels and downstream signalling of OLD-1. In addition, we also identify the transcription factor VAB-3, which is the *C. elegans* homolog of PAX6, to regulate the activation of the ORR through transcriptional regulation of *old-1.* This work describes a novel kinase-pseudokinase pair driving pathogen-specific immune signalling in the *C. elegans* epidermis that is spatially regulated by the activity of a conserved developmental factor.

## Results

### *old-1* and *flor-1* are required for oomycete recognition in *C. elegans*

To dissect the machinery involved in oomycete recognition, we performed a forward genetic screen using the induction of the *chil-27p::GFP* reporter as a read-out for pathogen detection. Animals carrying this marker were mutagenised with EMS and their F2 progeny was screened for loss of GFP expression upon addition of *M. humicola* extract. Whole genome sequencing of mutant animals identified multiple independent mutations in two genes, *old-1* and *flor-1* (Fig. 1A), which all led to complete loss of *chil-27p::GFP* induction upon extract treatment (Fig. 1B). Loss of *chil-27p::GFP* induction could be phenocopied by *old-1* and *flor-1* RNAi, and in a background carrying a genetic deletion of *old-1(ok1273),* suggesting that the recovered mutations represent loss-of-function alleles (Fig. 1B). OLD-1 and FLOR-1 are part of the KIN-16 family of receptor tyrosine kinases (RTKs), a family which is *C. elegans-*specific and its members are little characterised^16-18^.

**Figure 1.**
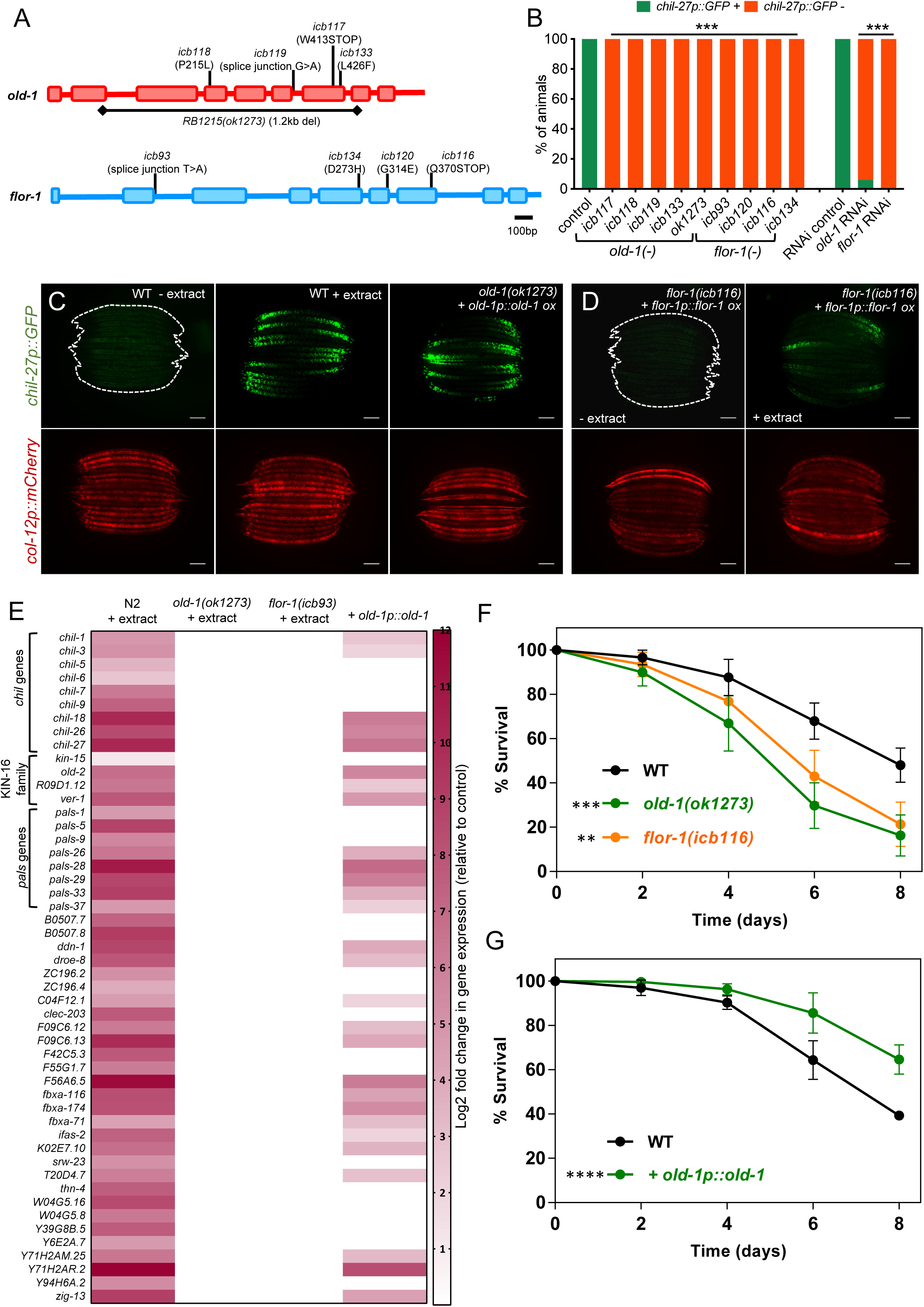
*old-1* and *flor-1* are required for oomycete recognition and immune response in *C. elegans*. **(A)** Gene structure of *old-1* and *flor-1* and location of mutations recovered through EMS screens that result in loss of *chil-27p::GFP* induction in response to *M. humicola* extract. **(B)** *icb117-119* contain loss of function mutations in *old-1*, *ok1273* contains a 1.2 kb deletion in *old-1*; all show a 100% loss of response to extract. *icb93, icb120* and *icb116* contain mutations in *flor-1* and also result in a complete loss of response to *M. humicola* extract. RNAi mediated knock-down of *old-1* or *flor-1* phenocopies the reduction in response to *M. humicola* extract (chi-square test for each sample, \*\*\**p* < 0.001, n > 50 for each sample). **(C)** Overexpression of *old-1(icbEx308)* in *old-1(ok1273)* causes constitutive activation of *chil-27p::GFP* mimicking that obtained upon extract treatment. **(D)** Overexpression of *flor-1(icbEx435)* in *flor-1(icb116)* mutant does not cause constitutive activation of *chil-27p::GFP* but rescues response to extract treatment. For C-D, presence of *col-12p::mCherry* represents a constitutive marker expressed in adults, n > 100 and scale bar is 100 µm. **(E)** Heatmap showing the induction of 50 most commonly upregulated ORR genes. Note that none of these ORR genes are transcriptionally upregulated in *old-1(ok1273)* and *flor-1(icb93)* mutants whereas around two-thirds are induced upon *old-1* overexpression (*icbEx308)* (compared to non-transgenic animals from the same population as control). Significant changes (*q* value < 0.1, *p* value < 0.01) are shown, white represents non-significant. Complete RNAseq tables are reported in Supplementary Table 1. **(F)** Loss-of-function of *old-1* or *flor-1* increase susceptibility to *M. humicola* infection. **(G)** Overexpression of *old-1p::old-1(icbEx358)* makes animals less susceptible to *M. humicola* infection. WT control used for this experiment also carries the hygromycin resistance (*icbEx360*). For panels F-G, n = 80-90 per condition, representative graph of three repeats shown, stars indicate significance with a Log rank test as follows: \*\**p* < 0.01, \*\*\**p* < 0.001, \*\*\*\**p* < 0.0001.

Overexpression of RTKs can lead to ligand-independent activation of downstream signalling events^19,20^ so we tested the effect of overexpressing the two identified RTKs in the respective mutants. While overexpression of *old-1p::old-1* resulted in constitutive activation of *chil-27p::GFP* just like extract treatment (Fig.1C), overexpression of *flor-1p::flor-1* in *flor-1(icb116)* mutants only restored the response to *M. humicola* extract (Fig. 1D).

To determine whether the loss of *chil-27p::GFP* induction in *old-1(-)* and *flor-1(-*) mutants upon extract treatment is representative of a broader loss of induction in ORR genes, we performed RNA-sequencing (Table S1). The transcriptomic results of wild type extract-treated animals were first compared to the previously published ORR dataset obtained^12^ using comparable conditions (L4 stage and 4 hours post extract treatment). We found 50 genes that are consistently upregulated in response to *M. humicola* extract in multiple independent experiments (Fig. 1E). These genes include members of the *chil* gene family, members of the KIN-16 family to which *old-1* and *flor-1* belong, and *pals* genes, which are also part of the intracellular pathogen response (IPR) induced upon infection with microsporidia or the Orsay virus in *C. elegans*^21^. We found that loss of *old-1* or *flor-1* function impaired the induction of all 50 genes upon extract treatment (Fig. 1E, Table S1). In contrast, more than 50% of these genes were found to be upregulated upon overexpression of *old-1* under its own promoter, suggesting that overexpression of *old-1* largely mimics the induction of ORR upon extract treatment (Fig. 1E, Table S1). Consequently, inability to activate the ORR made *old-1(-)* and *flor-1*(*-)* mutants survive less to infection by *M. humicola* than wild type animals (Fig. 1F); while constitutive activation of ORR through overexpression of *old-1p::old-1* caused increased survival to *M. humicola* infection (Fig. 1G). These results establish the requirement of OLD-1 and FLOR-1 as key players for the activation of oomycete recognition response in *C. elegans*.

### OLD-1 and FLOR-1 are required in the epidermis for the induction of ORR

To determine in which tissue *old-1* and *flor-1* are expressed, we performed single-molecule FISH (smFISH). For both genes, we observed epidermal expression as evidenced by the co-localisation of mRNAs with epidermal GFP-labelled nuclei (Fig. 2A). Interestingly, despite the *C. elegans* epidermis being a multinucleated syncytium^22^, *old-1* was found to be expressed at low levels exclusively in the anterior end of the epidermis, while *flor-1* mRNAs were more abundantly distributed throughout the epidermis in wild type animals (Fig. 2A). Translational reporters created by the fusion of OLD-1 and FLOR-1 with GFP also matched the mRNA expression and showed epidermal membrane localisation (Fig. 2B). Co-expression of OLD-1::GFP and FLOR-1::mScarlet showed co-localisation in the anterior epidermal membrane (Fig. S1A). Since both OLD-1 and FLOR-1 co-localise in the epidermal membrane, we wished to determine if they could be physically interacting to facilitate signalling. To this end, we used Fluorescent Resonance Energy Transfer (FRET) and found OLD-1::GFP and FLOR-1::mScarlet in close proximity to allow interaction both in the presence and absence of *M. humicola* extract (Fig. S1B). Finally, we tested whether epidermis-specific RNAi targeting *old-1* or *flor-1* would be sufficient to prevent induction of *chil-27p::GFP* upon extract treatment. We found that epidermal RNAi was sufficient to decrease *chil-27p::GFP* induction to the same extent as non-tissue-specific RNAi (Fig. 2C). Taken together, we conclude that these RTKs are likely to act together at the level of the epidermal membrane.

**Figure 2.**
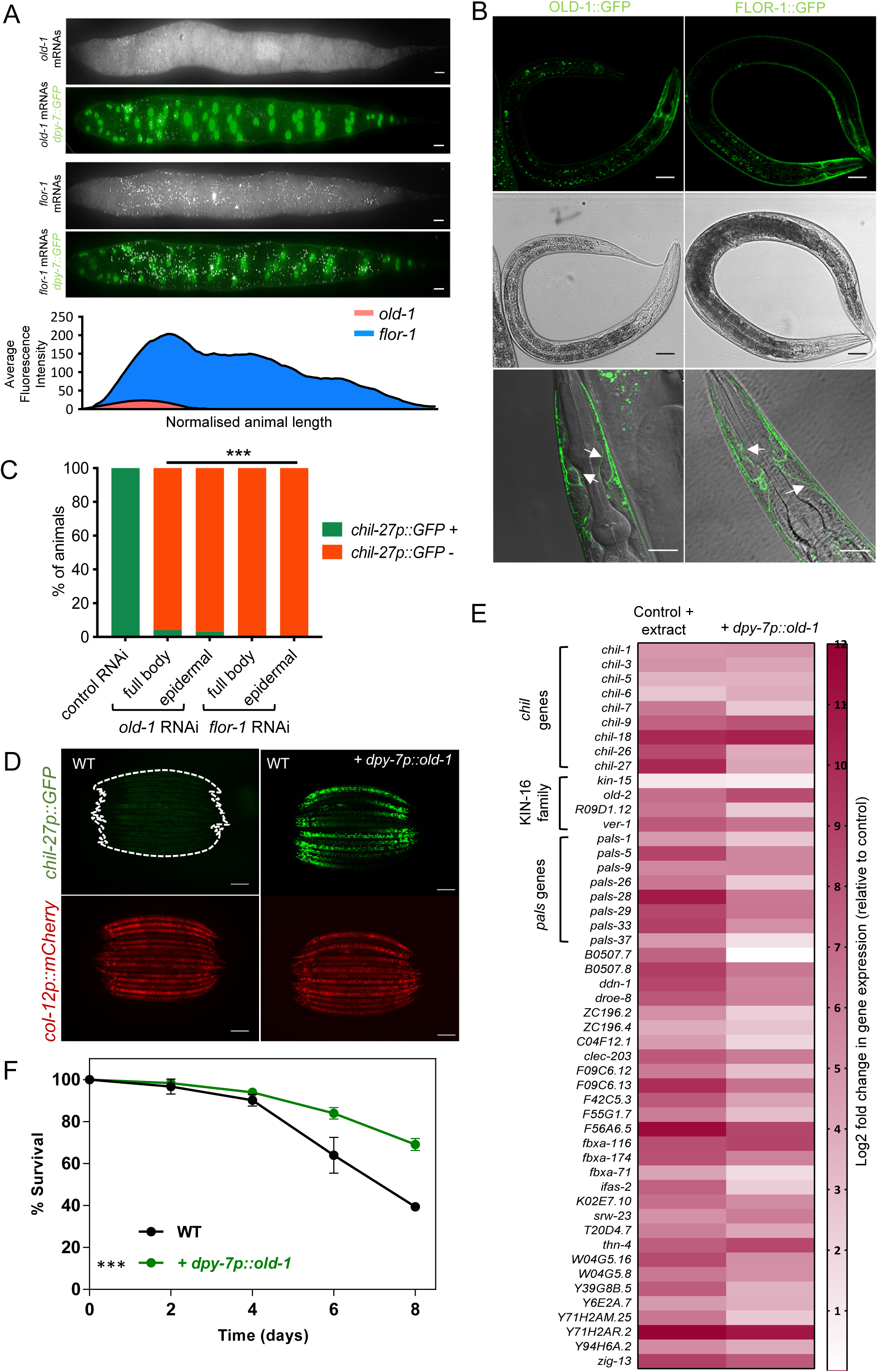
OLD-1 and FLOR-1 are required in the epidermis for the upregulation of ORR genes. **(A)** Expression of *old-1* and *flor-1* by smFISH. *old-1* and *flor-1* mRNAs (white spots) co-localise with epidermal nuclei (green) marked by a *dpy-7p::GFP* transgene. Note that *old-1* mRNAs are detected at the anterior side of the epidermis while *flor-1* mRNAs are detected throughout the epidermis. Scale bar is 10 µm. Average fluorescent intensity of mRNA signal for *old-1* and *flor-1* is plotted along a normalised body length. *flor-1* is expressed at a higher level throughout the epidermis whereas *old-1* is expressed at low levels in the anterior, n > 15 for each genotype. **(B)** An OLD-1::GFP *(icbEx310)* translational fusion is expressed primarily in the anterior epidermis, whereas a FLOR-1::GFP translational fusion *(icb136)* is expressed throughout the epidermis. Both translational fusions localise to the epidermal membrane (white arrows). Scale bar for top four images is 50 µm, and for bottom images is 20 µm. **(C)** Epidermal-specific RNAi knock-down of *old-1* and *flor-1* expression is sufficient to reduce response to *M. humicola* extract to the same level as whole-body RNAi knock-down, n > 50 for each treatment. Stars represent significance with a chi-square test, \*\*\**p* < 0.001. **(D)** Overexpression of *old-1* under an epidermal specific promoter (*dpy-7p*)(*icbEx309*) triggers constitutive activation of *chil-27p::GFP*. Note that expression is observed throughout the epidermis as opposed to the gradient seen upon endogenous overexpression and extract treatment, n > 100 animals, scale bar is 100 µm. **(E)** Heatmap showing relative expression of commonly upregulated ORR genes upon epidermal overexpression of *old-1(icbEx359)* and extract treated WT animals carrying hygromycin resistance (*icbEx360*). > 90% of these ORR genes were upregulated in an epidermal-specific *old-1* overexpression(*dpy-7p::old-1).* Significant changes (*q* value < 0.1, *p* value < 0.01) are shown, white represents non-significant. Complete RNAseq data are reported in Supplementary Table 2. **(F)** Representative survival assay for epidermal overexpression of *old-1*(*icbEx359*) which decreases susceptibility to infection by *M. humicola.* WT control used for this experiment also carries the hygromycin resistance (*icbEx360*) (n = 80-90 per condition, representative graph of three repats shown). Log rank test ***p < 0.001.

Next, we tested whether we could mimic the constitutive activation of *chil-27p::GFP* seen upon overexpression of *old-1*, using the *dpy-7* gene promoter that limits expression specifically to the epidermis. Here we found that *chil-27p::GFP* was expressed constitutively throughout the entirety of the epidermis (Fig. 2D, Fig. S2) as opposed to just the anterior side seen with extract treatment. This difference in *chil-27p::GFP* expression is consistent with the *old-1* mRNA distribution observed in the anterior side of the transgenics carrying *old-1* under its endogenous promoter, or throughout the epidermis when the *dpy-7* promoter was used instead (Fig. S2). RNAseq analysis of animals overexpressing *dpy-7p::old-1* showed upregulation of >95% of the key ORR genes (Fig. 2E, Table S2) Consequently, these animals also showed increased survival upon infection by *M. humicola* (Fig. 2F). As the activation of the ORR upon oomycete exposure is modulated by cross-tissue communication from neurons to epidermis^12^, we asked how overexpression of *old-1* relates to the requirement of neuronal signalling for activation of ORR. We have previously shown that *tax-4(ks11)* mutants show compromised induction of *chil-27p::GFP* upon treatment with diluted extract^12^. Here, we found that constitutive expression of *chil-27p::GFP* upon overexpression of *old-1* is not suppressed by *tax-4* loss of function, further suggesting that OLD-1 is likely to act at the level of the epidermis (Fig. S3).

### FLOR-1 is a pseudokinase that regulates OLD-1 membrane availability and its downstream signalling

Despite the fact that OLD-1 and FLOR-1 belong to the same RTK family, we did not see the same constitutive activation of *chil-27p::GFP* upon overexpression of *flor-1* (Fig. 1D). Upon closer investigation of key regions in the protein sequence that are known to be required for active kinase function, such as the glycine-rich, activation and catalytic loop^23,24^, we noticed key amino acid changes specifically in FLOR-1, which are likely to interfere with kinase activity (Fig. 3A). For example, a signature of catalytically deficient kinases, also known as pseudokinases, is mutations in an aspartate (broadly referred to as D166) and asparagine (N171), present within a HxDxxxxN motif required for catalysis. This motif was present in OLD-1 (<u>H</u>R<u>D</u>LALR<u>N</u>) and other family members, but not FLOR-1 (HR<u>A</u>LALR<u>S</u>). We used site directed mutagenesis to mutate these residues in the catalytic loop of OLD-1 to those found in FLOR-1 and changes to these amino acids resulted in the loss of constitutive activation of *chil-27p::GFP*, suggesting that an active kinase function is required for ligand independent downstream signalling upon overexpression of *old-1* (Fig. 3B). Unlike FLOR-1 overexpression under its endogenous promoter which does not spontaneously induce *chil-27p::GFP* (Fig. 1D), animals expressing high levels of FLOR-1 via the *dpy-7::flor-1* transgene show constitutive expression of *chil-27p::GFP* even in the absence of extract treatment (Fig. 3C). However, when the key catalytic residues (A297D, S302N, as present in OLD-1) are introduced back in FLOR-1, the induction of *chil-27p::GFP* is lost (Fig. 3C), while FLOR-1 retained the ability to rescue *chil-27p::GFP* induction in *flor-1(icb116)* mutants upon exposure to oomycete extract (Fig. 3C). These results indicate that FLOR-1 is likely to be a pseudokinase with reduced or no kinase activity.

**Figure 3.**
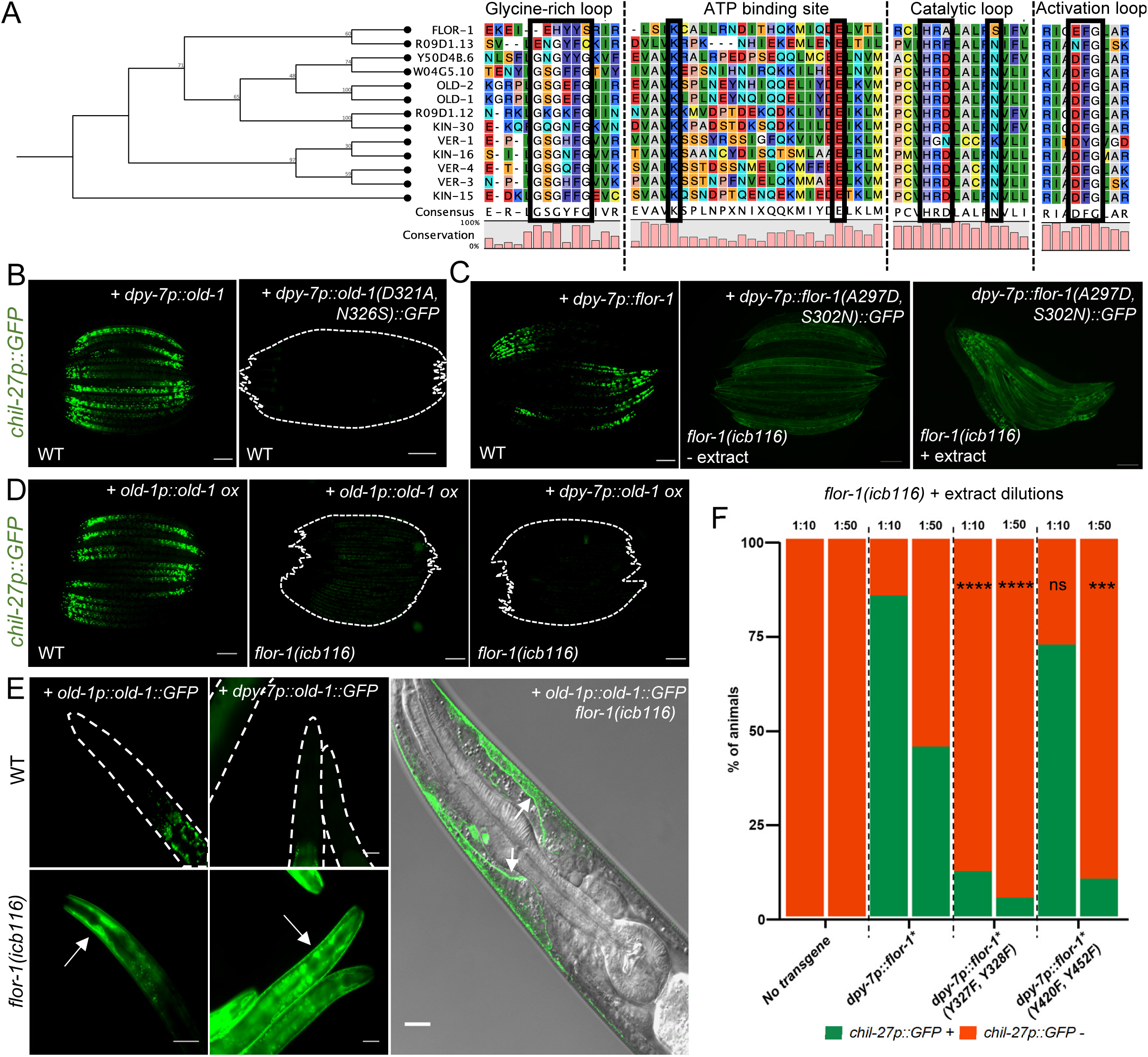
OLD-1 and FLOR-1 form a kinase-pseudokinase pair in the epidermal membrane. **(A)** Phylogenetic tree of the KIN-16 family of receptor tyrosine kinases focusing on alignment of key regions within the kinase domain. FLOR-1 lacks conserved kinase domain features, such as the conserved amino acids in the glycine loop (GxGxxGx), catalytic loop (HxDxxxxN), and activation loop (DFG motif) indicating that FLOR-1 is likely a pseudokinase. K72 salt-bridge and ATP-binding residue are present but the VAVK motif appears divergent in FLOR-1. Bootstrap values are shown at nodes of tree. Protein sequences were aligned, and phylogenetic tree was created using CLC Sequence Viewer (Qiagen). **(B)** Site directed mutagenesis of the catalytic loop of OLD-1 to the non-conserved amino acids found in FLOR-1 (D321A, N326S) inhibits upregulation of *chil-27p::GFP* upon epidermal overexpression *(icbEx316)*. **(C)** Site directed mutagenesis of the catalytic loop of FLOR-1*(icbEx426)* to the conserved amino acids found in OLD-1 (A297D, S302N) inhibits upregulation of *chil-27p::GFP* seen with *dpy-7p::flor-1(icbEx412)* but can still rescue response to extract in *flor-1(icb116)* mutants. **(D)** Constitutive activation of *chil-27p::GFP* upon overexpression of *old-1p::old-1 (icbEx433)* and *dpy-7p::old-1 (icbEx434)* is dependent upon functional FLOR-1. For B-D, n > 100 and scale bar is 100 µm. **(E)** Expression of *OLD-1::GFP (icbEx414, icbEx436*) is increased in the absence of *flor-1*. OLD-1::GFP accumulates at the epidermal membrane (white arrows) in *flor-1(icb116)* mutant. n > 100 animals, scale bar is 50 µm for left panels, 10 µm for right image. **(F)** Response to extract (dilutions 1:10 & 1:50) quantified in *flor-1(icb116)* mutants overexpressing different mutant forms of phosphorylation sites in *flor-1* in the hypodermis (n = 50 per strain, \*\*\*\**p* < 0.0001, \*\*\**p* < 0.001 with chi-square test). Phosphorylation mutants were created in *flor-1* sequence which was already carrying A297D and S320N mutations (*flor-1\**).

Next, we sought to determine the relationship between the two RTKs. To this end, we expressed *old-1* both under its endogenous promoter and under the epidermal-specific *dpy-7* promoter in a *flor-1(icb116)* mutant. In both cases, the constitutive expression of *chil-27p::GFP* was no longer observed (Fig. 3D), suggesting that FLOR-1 is required for OLD-1-mediated downstream signalling. To determine what happens to the expression and localisation of OLD-1 in the absence of *flor-1*, we expressed OLD-1::GFP in a *flor-1(icb116)* mutant. Unlike FLOR-1, the overall expression of OLD-1 fusion proteins observed in wild-type animals is low, even upon transgene overexpression at high concentrations with strong epidermal promoters (*dpy-7p*) and as part of multi-copy arrays (Fig. S4A). We hypothesised that low OLD-1 levels might reflect tight regulation to limit aberrant activation of downstream signalling. In fact, addition of the proteasome inhibitor bortezomib (BTZ) caused an increase of OLD-1 expression (Fig. S4B), supporting the hypothesis that excess OLD-1 is likely to be degraded by the proteasome. Surprisingly, in *flor-1(icb116)* mutants, we saw high levels of GFP expression with correct membrane localisation in the anterior epidermis when OLD-1 was expressed under its endogenous promoter (*old-1p::old-1::GFP*), and expression throughout the epidermis when OLD-1 was expressed under the epidermis-specific promoter (*dpy-7p::old-1::GFP)* (Fig. 3E). Additionally, an independent EMS suppressor screen performed on animals constitutively expressing OLD-1::GFP *(icbIs22)* identified five further alleles of *flor-1,* which fully suppressed both constitutive activation of *chil-27p::GFP* (data not shown) and also caused accumulation of OLD-1::GFP in the anterior epidermis (Fig. S4C). These results substantiate the requirement of FLOR-1 for regulating the downstream signalling pathway and OLD-1 levels at the epidermal membrane.

Given that FLOR-1 appeared to act downstream of OLD-1, we next asked if OLD-1 could activate the downstream signalling pathway by phosphorylating FLOR-1. To answer this, we mutated specific tyrosine residues in FLOR-1 to phenylalanine, and tested their ability to rescue *chil-27p::GFP* induction in *flor-1(icb116)* mutants upon oomycete extract treatment. We chose to mutate the two adjacent tyrosines within the activation loop (Y327, Y328), and two tyrosines (Y420 and Y452) towards the C-terminal end of FLOR-1. The rationale was that activation loop tyrosine residues have been previously shown to be important for the function of pseudokinases^25^; and that a *flor-1(icb126)* allele recovered from our OLD-1::GFP suppressor screen, which corresponds to a truncated protein lacking tyrosines Y420 and Y452. We used FLOR-1*(A297D, S302N)* as the template for mutagenesis because its overexpression does not induce *chil-27p::GFP* constitutively. *flor-1(icb116)* mutant animals carrying the respective constructs were treated with oomycete extract at the early L1 stage and induction of *chil-27p::GFP* was scored after 24 hours. While FLOR-1 overexpression was strongly able to rescue *chil-27p::GFP* induction in *flor-1(icb116)* upon extract treatment, the response obtained with FLOR-1(Y327F, Y328F) and to a lesser extent with FLOR-1 (Y420F, Y452F) was significantly reduced upon extract (Fig. 3F). These results suggest that OLD-1 mediated phosphorylation of multiple tyrosine residues in FLOR-1 is required for the activation of downstream signalling pathway, with the phosphorylation of tyrosine residues in the activation loop being more critical than the others.

### The transcription factor VAB-3 transcriptionally regulates *old-1* to activate the ORR

In the screen mentioned above aimed at recovering suppressors of the *old-1* overexpression phenotype, we also obtained a missense mutation (*icb127*), which changed an R to Q amino acid within the homeodomain of the transcription factor VAB-3. VAB-3 is the *C. elegans* PAX6 homologue and is well-studied for its role in head morphogenesis and cell fate specification in anterior epidermal cells^26,27^. The *icb127* mutation caused loss of constitutive *chil-27p::GFP* expression without accumulation of OLD-1::GFP as previously observed in *flor-1(icb116)* mutants (Fig. 4A). In addition, this mutant failed to induce *chil-27p::GFP* upon exposure to oomycete extract (Fig. 4A). To confirm that *icb127* is a new loss-of-function allele of *vab-3*, we obtained animals with another previously reported loss-of-function allele *vab-3(ju468)* and crossed them with the *chil-27p::GFP* reporter. We found that the *vab-3(ju468)* mutation suppressed *chil-27p::GFP* induction both upon extract treatment as well as upon overexpression of *old-1* (Fig. 4B). Overexpression of WT *vab-3* through a fosmid rescued constitutive *chil-27p::GFP* expression upon *old-1* overexpression in *vab-3(icb127)* (Fig. 4C). These results suggest that *vab-3* is a key regulator of the response to oomycete recognition.

**Figure 4.**
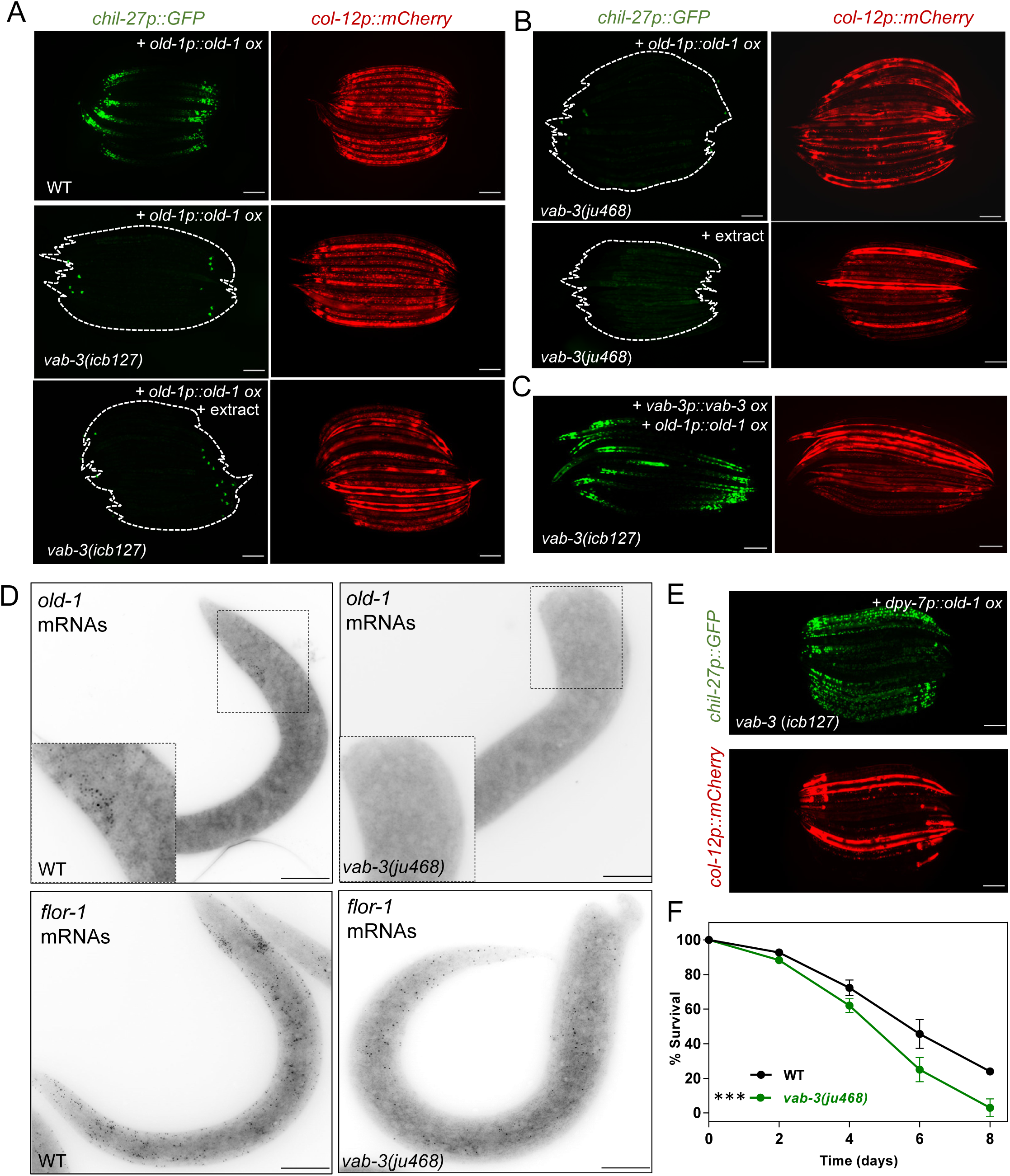
PAX-6 homolog VAB-3 regulates *old-1* expression. **(A)** A mutation in *vab-3(icb127)* inhibits constitutive activation of *chil-27p::GFP* and induction of *chil-27p::GFP* in response to *M. humicola* extract. **(B)** This effect was phenocopied by an independent *vab-3* mutant allele (*ju468).* **(C)** Constitutive expression of *chil-27p::GFP* was restored in *vab-3(icb127)* carrying an *old-1p::old-1* transgene by overexpressing a *vab-3* containing fosmid (WRM0640cC05).**(D)** A loss of *old-1* mRNA expression is seen in in *vab-3(ju468),* but *flor-1* mRNAs remain unaffected. For D, scale bar is 10 µm and n > 15. **(E)** Constitutive activation of *chil-27p::GFP* is observed in *vab-3(icb127)* upon overexpression of *dpy-7p::old-1*. For A-C & E, n >100 animals and scale bar is 100 µm, presence of the *chil-27p::GFP* reporter is indicated by *col-12p::mCherry* also present in the same transgene. **(F)** *vab-3(ju468)* mutants are hypersusceptible to infection by *M. humicola* (n = 90 per condition, representative graph of three repeats shown) *** *p* < 0.001 Log-Rank test.

Using smFISH, we found localisation of *vab-3* mRNAs in the anterior epidermis as expected^26^ (Fig. S5). Since this pattern is reminiscent of *old-1* expression (Fig. 2A), we asked if VAB-3 could be regulating *old-1* expression. Indeed, we did not detect any *old-1* mRNAs in *vab-3(ju468)*, while *flor-1* levels and localisation were not affected (Fig. 4D). This means that lack of *chil-27p::GFP* induction in *vab-3* mutants is likely a consequence of loss of *old-1* expression. To test this hypothesis, we overexpressed *old-1* in *vab-3(icb127)* under an epidermis-specific *dpy-7* promoter to circumvent the requirement for VAB-3 regulation and found that the *vab-3* mutation no longer suppressed the constitutive *chil-27p::GFP* expression (Fig. 4E). Finally, just like *old-1(ok1273)* mutants we found *vab-3(ju468)* to be hypersusceptible to infection by *M. humicola* (Fig. 4F). To conclude, these results allow us to propose a model where OLD-1 is an epidermal RTK whose expression is transcriptionally regulated by VAB-3 and membrane localisation by another RTK family member FLOR-1. When OLD-1 is activated upon oomycete exposure or overexpression, it can phosphorylate FLOR-1 and trigger the activation of ORR mediating oomycete resistance (Fig. 5).

**Figure 5.**
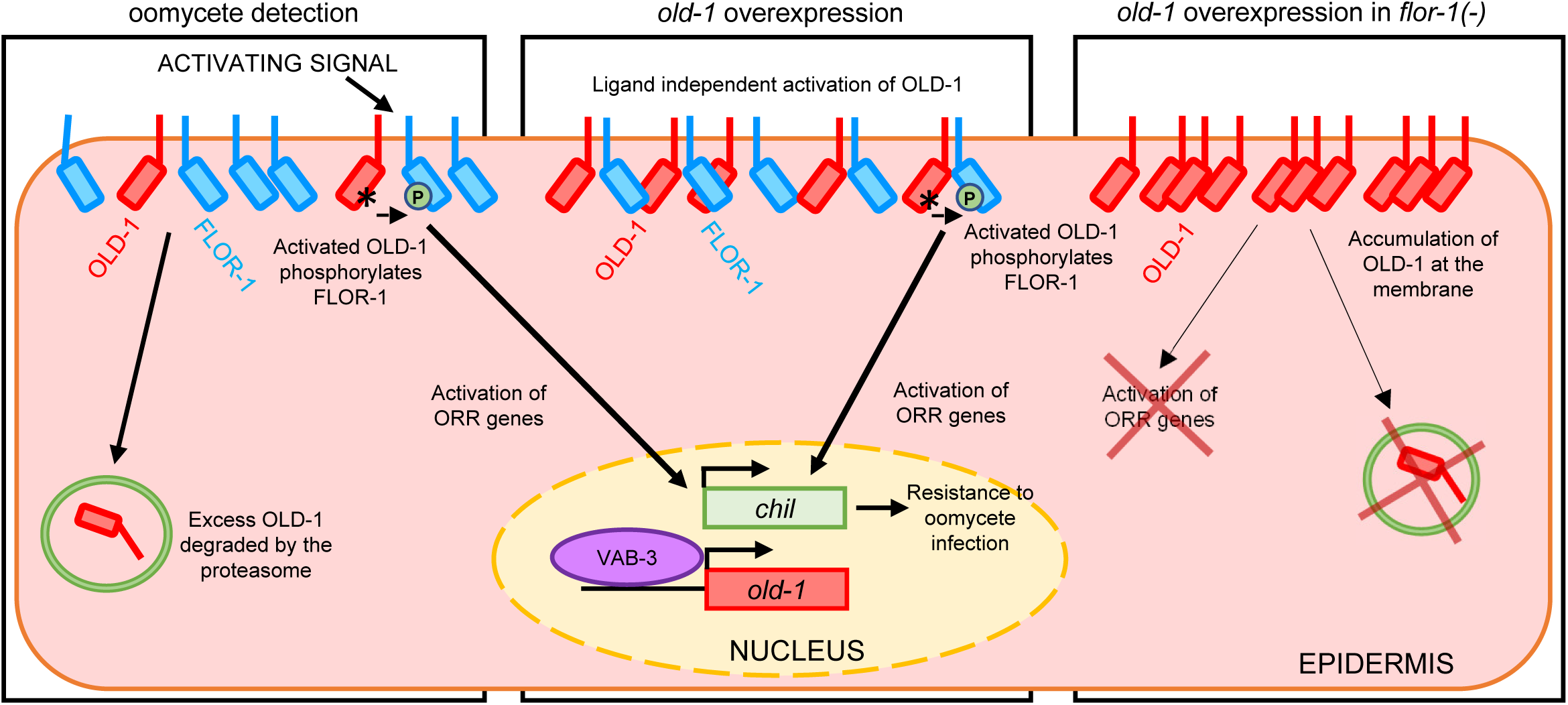
Schematic model for signalling through OLD-1/FLOR-1. Schematic model of the possible interaction between OLD-1 and FLOR-1 at the epidermal membrane upon oomycete detection, and *old-1* overexpression in a WT or *flor-1(-)* mutant background. FLOR-1 is both required to facilitate signalling of the oomycete recognition response through phosphorylation by OLD-1 and for maintaining low levels of OLD-1 at the epidermal membrane. VAB-3 is a transcription factor that acts to regulate *old-1* expression in the context of the response to extract and the constitutive activation of *chil-27p::GFP* upon *old-1* overexpression.

## Discussion

Building upon our previous discovery of cross-tissue signalling in the oomycete recognition response in *C. elegans*^12^, we present here two key regulators of this program namely OLD-1 and FLOR-1. These are members of the KIN-16 family of receptor tyrosine kinases^17^ and are necessary to drive the epidermal component of the defence response against oomycete infection. While *old-1* has been previously implicated in lifespan extension and stress resistance^28^, there is no current link between KIN-16 family members and immunity. However, several members of this family, such as *kin-15*, *old-2, ver-1* and *R09D1.12* are induced as part of the ORR and thus may contribute to immunity. It is of note that both *old-1* and *flor-1* expression is not induced upon infection, so these new regulators could not have been identified through transcriptomic profiling. This is similar to other gene families like the *pals* genes, where some genes are part of the IPR and others regulate the IPR. It is also interesting that the *old-1* locus as well as several other members of this family, such as *kin-15, kin-16, R09D1.12, old-2* and *R09D1.13,* are located within the previously identified *chil* gene clusters on chromosome II^8^. Genomic clustering of *C. elegans* immune response genes has been reported previously as well, for example on chromosome IV, which contains genes induced in response to *Microbacterium nematophilum*^29^, the *nlp* and *cnc* gene clusters upregulated in response to the fungus *Drechmeria coniospora*^30^, or the *pals* gene cluster on chromosome I that contains 13 *pals* members. Since *old-1* regulates *chil* gene expression to antagonise infection by *M. humicola,* our study suggests that the close proximity of KIN-16 family members and *chil* genes might be indicative of a cluster of resistance genes acting together as part of the signalling pathway underlying oomycete-specific immunity.

Alignment of all members of the KIN-16 family revealed that FLOR-1 contains key amino acid changes in highly conserved regions shown to be required for full kinase activity. 10% of human RTKs are thought to be missing at least one conserved, catalytically important residue and are thus classified as putative pseudokinases^23^. First, the glycine loop with the consensus sequence GxGxxG, which plays a role in phosphate binding, is missing in FLOR-1. Substitution of the first two glycines is thought to distort the ATP binding site^24^. Second, the catalytic loop of RTKs contains two conserved residues – the aspartate in the HRD motif is thought to function as the catalytic base for phosphorylation and an asparagine is required for metal ion binding for catalysis^24^. FLOR-1 does not have either of these conserved amino acids. Third, the VAIK motif co-ordinates the co-factor ATP, anchoring and orienting the ATP and forming a salt bridge with E91^31^. Although the K/E residues are present in FLOR-1, the motif does stray from the consensus in the family (LSIK vs VAVK) and may hinder ATP binding. Finally, the DFG motif of the activation loop has also diverged from the consensus in the family– the aspartate is thought to be one of the most critical amino acids for active kinase function as it is required for divalent cation binding required for catalysis^24^. FLOR-1 does not have the conserved aspartate in this motif, instead it has glutamate. Although glutamate and aspartate are chemically similar, it has been shown that this substitution is sufficient to inactivate phospho-transfer in kinases^32-34^. In summary, while a single change in any of these conserved motifs is usually sufficient for putative classification as a pseudokinase, FLOR-1 has changes in at least 4 out of these 6 critical amino acids. Nevertheless, FLOR-1 is functional and biologically important because it is required for immune signalling upon oomycete recognition acting downstream of the OLD-1 receptor tyrosine kinase.

The identification of FLOR-1 to play a role in oomycete resistance expands the repertoire of known pseudokinases that are linked with immune signalling in *C. elegans*. For example, the putative pseudokinase *nipi-3* acts as a negative regulator of transcription factor *cebp-1* and a positive regulator of transcription factor *skn-1* in the intestinal immune response against *Pseudomonas aeruginosa*^35,36^. *nipi-3* is also necessary for the *D. coniospora*-specific upregulation of *nlp-29* that triggers the p38-signalling cascade^37^, but the exact mechanisms are not well understood. *nipi-3* is part of the Tribbles pseudokinase family that have lost active kinase function, but act as scaffold or adaptor proteins and are involved in assembly or regulation of signalling components^38^. Part of the same family, *nipi-4* is a pseudokinase also involved in the positive regulation of *nlp* genes after *D. coniospora* infection, in addition to controlling the constitutive activation of antimicrobial *cnc* genes regulated by the TGF-β pathway^39^.

It is thought that some pseudokinases, despite their unusual sequence features, may still be able to catalyse phosphotransfer albeit to a limited extent^24,31,40^. For example, HER3 (human epidermal growth factor receptor 3, also known as ErbB3) is a key example of a low activity kinase. It is one of four members of the human epidermal growth factor receptor tyrosine kinase family, but stands out as the only member to have changes in the conserved ATP binding region^41^. This family is well studied due to its implication in tumorigenesis and is also a target for several anti-cancer drugs^42^. HER3 lacks the glutamate in the ATP binding region (substituted for a histidine) and replaces the aspartate in the HRD motif of the catalytic loop with an asparagine, leading to the initial hypothesis positing that HER3 is unable to bind ATP and is thus catalytically inactive^43^. However, follow up studies have shown that HER3 is in fact able to bind ATP and capable of trans-autophosphorylation^44,45^. Interestingly, this activity is much lower than other HER proteins and is dependent upon its active kinase partner, HER2^44,46^. The weak kinase activity may be sufficient to trans-phosphorylate within a HER2/HER3 heterodimer, activating the “active” kinase partner which then uses its increased kinase capabilities to phosphorylate exogenous substrates^44^. While we cannot rule out that FLOR-1 may have some weak kinase activity, it appears that it is phosphorylated downstream of OLD-1. Therefore, FLOR-1 is more likely to serve as a scaffold for assembly of signalling proteins leading to activation of ORR. In future, phosphoproteomic-based identification of OLD-1 targets, coupled with further genetic and RNAi screens can help elucidate the tissue-specific signalling network involved downstream of OLD-1/FLOR-1 in responding to oomycete recognition in *C. elegans*.

Our study also provides evidence for a link between the homeobox transcription factor VAB-3/PAX6 and the regulation of immune signalling in *C. elegans*. PAX6 is a highly conserved transcription factor, critical for its role in visual system development, and mutations in PAX6 are known to be associated with a wide array of symptoms, most notably congenital eye malformations^47,48^. An intestinal homeobox gene in *Drosophila* has been shown to repress nuclear factor kappa B-dependent antimicrobial peptide expression^49^. Our results also suggest that a conserved developmental regulator of anterior epidermal morphogenesis also regulates the expression of a signalling component that is required for the oomycete recognition response. Interestingly, this regulation can shape in this case the spatial pattern of immune signalling activation and readily explains the characteristic gradient of *chil-27* induction from the anterior epidermis to the posterior body upon pathogen recognition. The core sequence (5’-GCGTA-3’) associated with VAB-3 binding^50,51^ is present upstream of the *old-1* transcription start site, suggesting that VAB-3 is likely to regulate the transcription of *old-1* in a direct manner. It would be interesting to see in the future whether PAX6 is required for shaping immune responses to infection by oomycetes or other natural pathogens in different animals.

## Methods

### C. elegans and M. humicola maintenance

All *C. elegans* strains were cultured and maintained on NGM plates seeded with *E. coli* OP50 under standard conditions^52^. *M. humicola* was maintained by chunking infected worms onto fresh NGM plates and supplementing with fresh N2. For use in experiments, dead animals filled with sporangia were picked and transferred to the *E. coli* lawn of fresh NGM plates.

### EMS mutagenesis

Animals carrying the *chil-27p::GFP* reporter (*icbIs5)* were mutagenised using the chemical mutagen ethyl methanesulfonate (Sigma). L4 stage animals (P0) were incubated with 24 mM EMS in 4 mL M9, for 4 hours. Worm pellets were washed 10 times with 15 mL M9 to remove residual EMS. F2s were treated with extract and animals no longer expressing the *chil-27p::GFP* reporter were selected. Similarly, for the *old-1p::old-1* suppressor screen, F2 animals (*icbIs22)* no longer constitutively expressing *chil-27p::GFP* were selected. Around 60,000 haploid genomes were screened in both cases to identify suppressor mutants. *old-1* and *flor-1* mutants were mapped by crossing to the highly polymorphic CB4856 strain, F2 recombinant genomic DNA was isolated and sent for whole genome sequencing, completed by Novogene (Cambridge). WGS data was analysed using the CloudMap Hawaiian variant mapping pipeline on a Galaxy Server to identify causative mutations^53,54^. *icb127* was mapped using the mapping-by-sequence pipeline MIModD (https://mimodd.readthedocs.io/en/latest/index.html). In brief, variant allele frequency was mapped by first aligning the mutant genome sequence to N2 reference genome, followed by a comparison with single nucleotide polymorphisms found in CB4856. Regions enriched for N2 were then analysed for candidates, which were homozygous mutations not found in the CB4856 background.

### Infection and induction assays

Infection assays were performed in triplicate at 20°C on NGM plates seeded with 100 µl OP50. Five *M. humicola* infected (dead) animals were added to the OP50 lawn of four plates and minimum of 20 L4s were transferred to each plate, meaning a minimum of 80 animals were used per biological repeat. Dead animals with visible sporangia were scored every 48 hours and live animals were transferred to a new NGM plate supplemented with an additional five *M. humicola* infected (dead) animals. Dead animals without evidence of infection or missing animals were censored from the counts. Representative graph of three biological replicates with a minimum of 80 animals per biological replicate are shown in figures.

*M. humicola* extract was made and induction assays were performed as described previously^12^. *M. humicola* extract was added to the OP50 lawn when *C. elegans* eggs were added to the plate or at L2 stage. Presence or absence of *chil-27p::GFP* was quantified at L4 or day-1 adult stage using a using a Zeiss Axio Zoom V16 microscope.

### RNA-sequencing

For all RNA-seq experiments, animals were synchronised via bleaching and collected in triplicate for RNA extraction at L4 stage. Where extract was used, 200 µl was added to the OP50 lawn 4 hours before animal collection. Total RNA was extracted using TRIzol (Invitrogen) and transcriptome sequencing was completed by BGI (Hongkong) and Novogene (Cambridge). Kallisto^55^ was used for alignments with the WS283 transcriptome from Wormbase. Count analysis was performed using Sleuth^56^ along with a Wald Test to calculate log2fold changes. All RNAseq data files are publicly available from the NCBI GEO database under the accession number GSE220958.

### Microscopy

Single molecule FISH sample were fixed and prepared for microscopy as described previously^57^. In brief, animals were bleached and fixed at L2 stage using 4% formaldehyde (Sigma-Aldrich) for 45 mins and permeabilised with ethanol for at least 24 hours. Samples were then hybridised with a probe for 16 hours (probe sequences included in Table S3). Imaging was performed using the 100x objective of a Nikon Ti-eclipse inverted microscope fitted with an iKon M DU-934 CCD camera (Andor) and operated via the Nikon NIS Elements Software. Images were analysed using ImageJ FIJI software.

Confocal microscopy was performed on a Leica SP8 inverted Confocal Microscope with animals anaesthetised using 10 mM sodium azide and mounted on a 2% agarose pad.

Prior to FRET imaging, mixed stage animals were fixed using 4% formaldehyde (Sigma-Aldrich) for 45 mins and mounted on a 2% agarose pad. Where extract was used, 200 µl was added to the plate 24 hours before fixation. Microscopy was performed using a Leica Stellaris 5 in Confocal mode to perform acceptor depletion FRET at 63x magnification. GFP and mScarlet expression was measured three times prior to bleaching using a 488 nm and 561.4 nm laser respectively. mScarlet was then photobleached using the 561.4 nm laser and GFP and mScarlet expression were measured three times post photobleaching. Change in fluorescent intensity was analysed using FIJI macros developed by the Imperial College FILM facility.

### Molecular cloning and transgenesis

For rescue of mutant phenotypes and initial overexpression of *old-1*, gene sequences were PCR amplified from N2 genomic DNA using primers old-1FullF and old-1FullR and injected at 20 ng/µl. *old-1::GFP* was PCR amplified from the fosmid CBGtg9050E024D (TransgeneOME) using primers old-1FullF and old-1FullR and injected at 50 ng/µl. For rescue of *flor-1(icb116)*, the full gene locus was amplified from genomic DNA using primers T01G5.1promF and T01G5.1FullR2 and injected at 20 ng/µl. *flor-1::flor-1::GFP* was PCR amplified from MBA998 using the primers T01G5.1promF and T01G5.1FullR2 and injected at 30 ng/µl. The *flor-1::flor-1::mScarlet* plasmid (pER6) was created using Gibson cloning into a pER4 *(wrmScarlet::unc-54 3’UTR)* backbone using primers T01G5.1promF and T01G5.1R_pER2/4 to amplify the gene locus and injected at 40 ng/µl.

All constructs expressed under the *dpy-7* promoter were inserted into plasmid pIR6 digested with FastDigest enzyme SmiI, using Gibson Cloning. The *dpy-7p::old-1* plasmid (pFD20) was created by amplifying *old-1* from gDNA using primers dpy-7old-1gibsonfwd and dpy-7old-1gibsonrev. The *dpy-7p::old-1(D321A, N326S)::GFP* plasmid (pFD23) was created using a Site Directed Mutagenesis Kit (Agilent) to introduce mutations into pFD21 using primers old-1kinasemutF and old-1kinasemutR. The *dpy-7p::flor-1::GFP* plasmid (pJS1) and the *dpy-7p::flor-1* plasmid (pJS2) were made using primers dpy-7T01G5.1gibF and T01G5.1unc-54 gibR to amplify *flor-1::GFP* and *flor-1* from MBA998 and gDNA respectively. Site directed mutagenesis was carried out on a *dpy-7p::flor-1::GFP* plasmid (pJS1) to introduce key OLD-1 catalytic residues back into FLOR-1. The resulting *dpy-7p::flor-1(A297D, S302N)::GFP* plasmid (pJS4) was made using primers T01G5.1 SDM F and T01G5.1 SDM R. To mutate specific tyrosines in FLOR-1, site directed mutagenesis was carried out on the plasmid pJS4. Residues Y327 and Y328 in the FLOR-1 activation loop were mutated using primers FLOR-1 SDM AL F2 and FLOR-1 SDM AL R2 to generate the *dpy-7p::flor-1*(Y327F, Y328F)::GFP* plasmid (pJS6). Residue Y420 in FLOR-1 was mutated using primers T01G5.1 Y420 SDM F and T01G5.1 Y420 SDM R. Residue Y452 in FLOR-1 was subsequently mutated using primers T01G5.1 Y452 SDM F and T01G5.1 Y452 SDM R to generate the *dpy-7p::flor-1*(Y420F, Y452F)::GFP* plasmid (pJS7). All constructs under the *dpy-7* promoter were injected at 5 ng/µl.

The endogenous copy of *flor-1* was tagged with codon optimised GFP (GFPo) at the C-terminus using a CRISPR/Cas-9 GFP approach. GFPo was amplified from plasmid pDK57 using two sets of primers: GFPoATG-F and GFPoSTOP-R and T01G5.1Frep and T01G5.1Rrep, which carry 120 bp homology either side of the Cas-9 cut site. Equimolar quantities of these two PCR products were melted and reannealed to create a repair template with a single-stranded homology on each arm. The repair template was injected at 0.11 ng/µl into WT animals along with 0.25 ng/µl protein Cas-9 (IDT), 0.02 ng/µl tracrRNA (IDT), 0.02 ng/µl crRNA (IDT) and 40 ng/µl *rol-6 (su1006)* ^58^.

Roller animals were selected and genotyped using primers T01G5.1fwd and GFPoSTOPR. The repair template introduced 3 synonymous changes at the end of the *flor-1* coding sequence to prevent re-cutting of the gene. The *icb127* mutant was rescued using injection of fosmid WRM0640cC05(Source BioScience) at 10 ng/ul.

### RNAi

RNAi experiments were performed using the feeding method with clones obtained from the Ahringer Library (Source BioScience). Expression of dsRNA was induced by the addition on 1 mM IPTG (Sigma) to NGM plates. 6 L4s were added to RNAi plates and scored 72 hours later, or bleached eggs were added and scored 48 hours later, for *chil-27p::GFP* induction. Where extract was used, 200 µl was added 48 hours before scoring.

### BTZ treatment

L2 animals were treated with different concentrations of Bortezomib (Sigma-Aldrich) for 24 hours prior to imaging. DMSO was used as a vehicle control.

## Acknowledgements

We thank Vladimir Lazetic and Spencer Gang for comments on the manuscript. Some strains were provided by the CGC, which is funded by NIH Office of Research Infrastructure Programs (P40 OD010440). We acknowledge the support from the Wellcome Trust [219448/Z/19/Z].

## Supplementary Figure Legends

**Figure S1.**
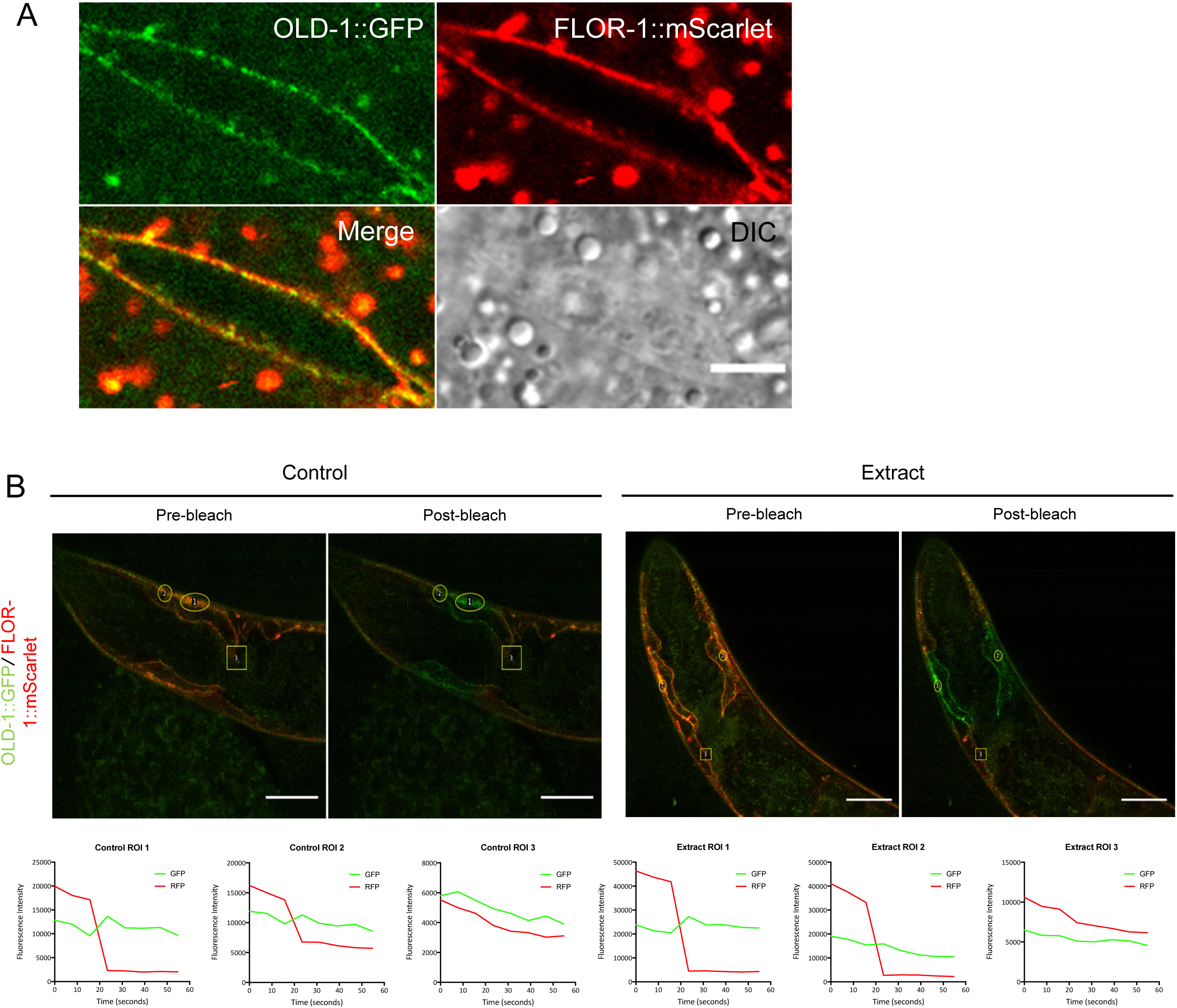
FRET shows that OLD-1 and FLOR-1 are in close enough proximity to be interacting at the epidermal membrane. **(A)** OLD-1::GFP and FLOR-1::mScarlet (*icbEx364*) co-localise in the anterior epidermis. Close-up of an anterior seam cell, scale bar is 5 µm. **(B)** Pre-bleach image shows co-localisation of OLD-1::GFP and FLOR-1::mScarlet in the anterior hypodermis. Yellow circle shows regions of interest (ROIs) targeted for bleaching, yellow square shows control region outside of bleached zone. Post-bleach image shows fluorescence expression after mScarlet has been photo bleached. Graphs show change in GFP and mScarlet fluorescence intensity at before and after photobleaching. An increase in GFP after mScarlet is bleached indicates that OLD-1::GFP and FLOR-1::mScarlet are within 10 nm of each other on the membrane and possibly interact. Both ROI1 and ROI2 show an increase in GFP expression after mScarlet is bleached both with and without extract. This increase is not seen in the negative control ROI3. Therefore, OLD-1 and FLOR-1 are found in close enough proximity to be interacting in the presence and absence of extract treatment. Representative image and graphs, n > 10 per treatment, scale bar is 10 µm.

**Figure S2.**
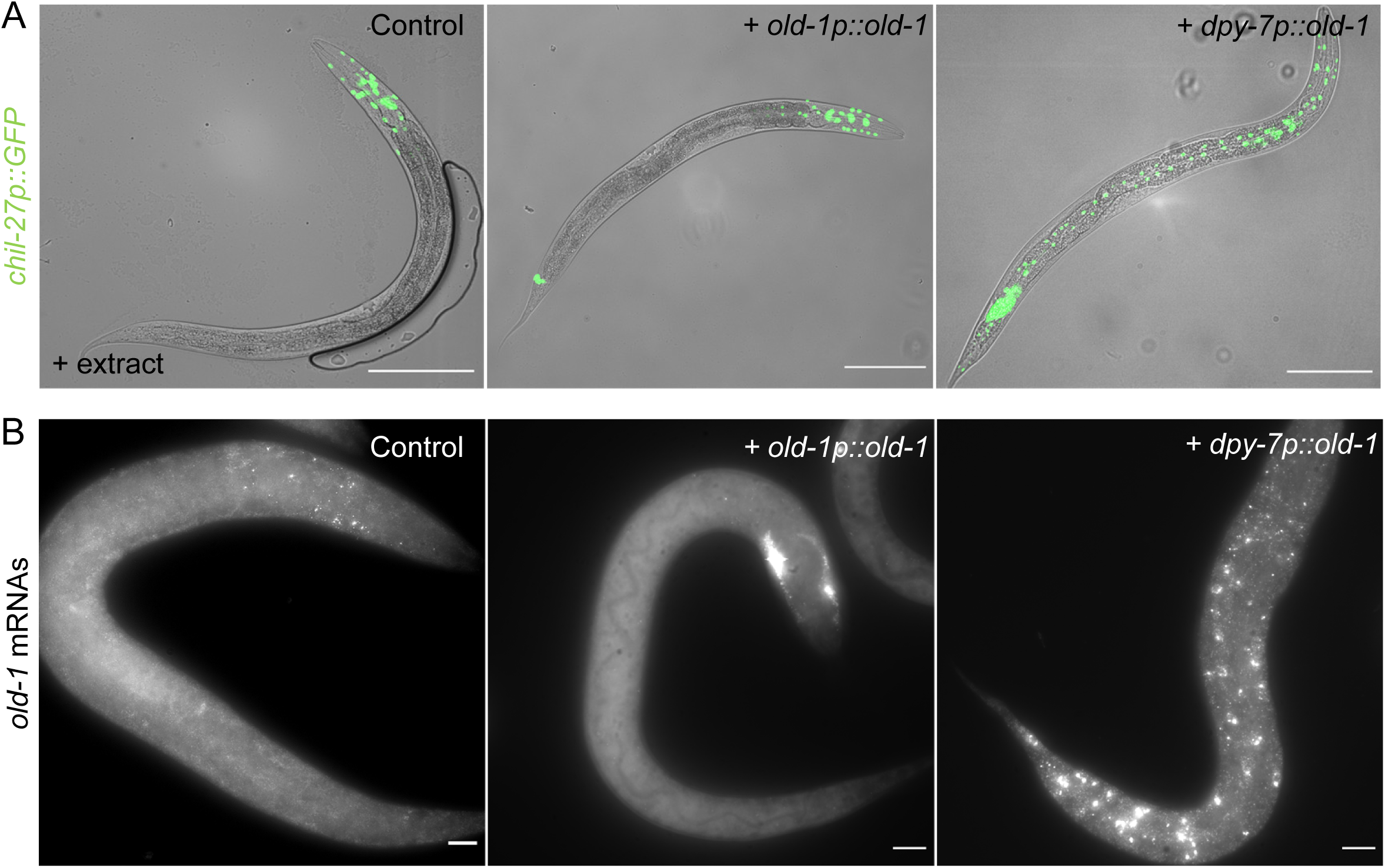
*dpy-7p::old-1* overexpression causes expansion of *old-1* mRNAs throughout the epidermis. **(A)** The *chil-27p::GFP* transcriptional reporter is expressed in an anterior biased gradient upon extract treatment and *old-1p::old-1* overexpression (*icbIs22).* This signal expands throughout the epidermis in *dpy-7p::old-1 (icbEx309)* animals. Scale bar is 100 µm, n > 100, *bus-1p::GFP* in the tail represents the co-injection marker. **(B)** In WT animals, *old-1* mRNAs are expressed in the anterior at low levels. *old-1p::old-1* overexpression (*icbEx358*) increases mRNAs in the anterior epidermis. *dpy-7p::old-1* overexpression (*icbEx359*) results in *old-1* mRNAs and transcription foci being present throughout the length of the body. Scale bar is 10 µm, n > 15.

**Figure S3:**
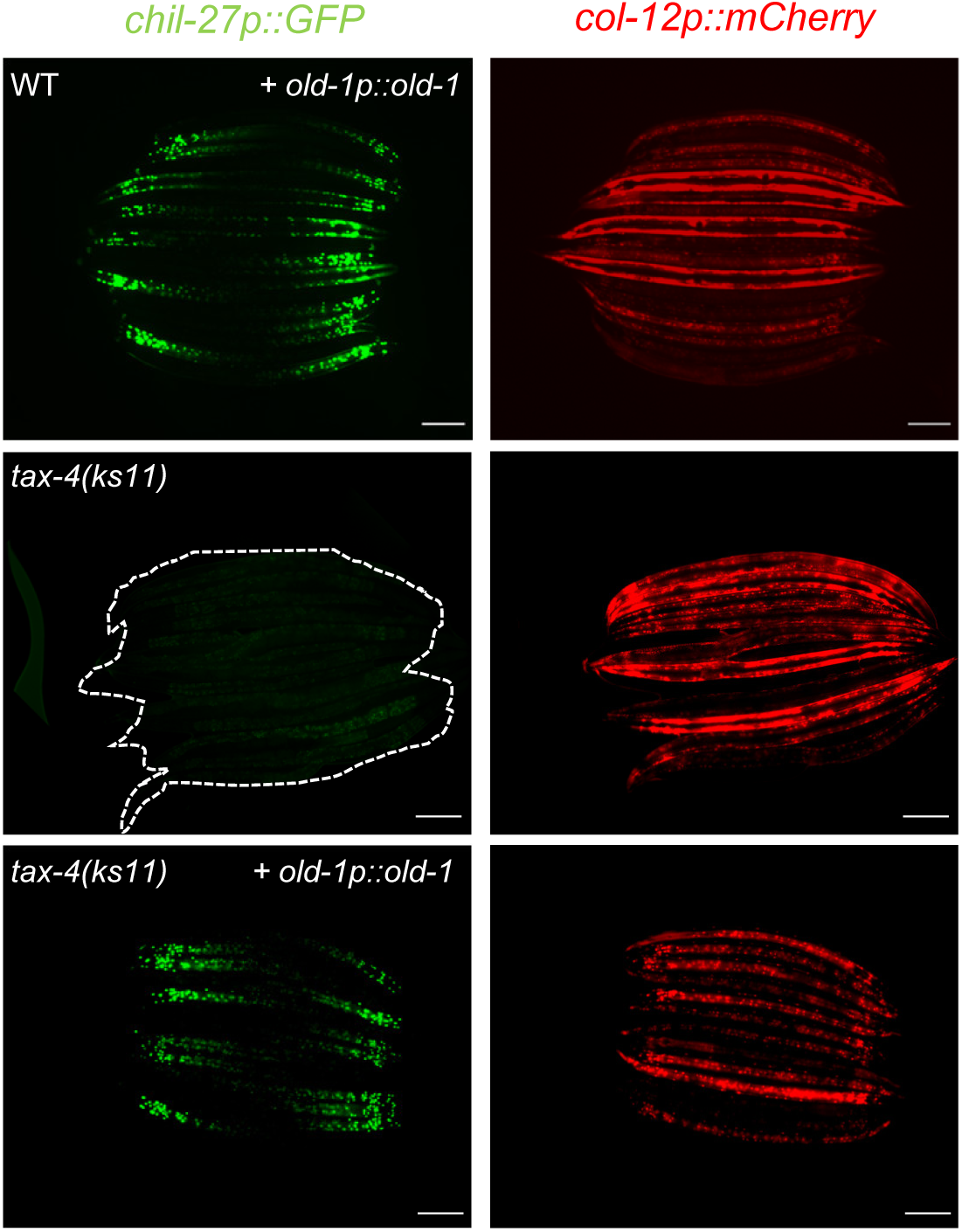
*old-1* overexpression does not require TAX-4-dependent signalling for induction of *chil-27p::GFP*. Loss of *tax-4(ks11)* function does not suppress the constitutive activation of *chil-27p::GFP* upon *old-1* overexpression (*icbEx424*), n > 100 animals, scale bar is 100 µm, presence of the *chil-27p::GFP* reporter is indicated by *col-12p::mCherry* also present in the same transgene.

**Figure S4.**
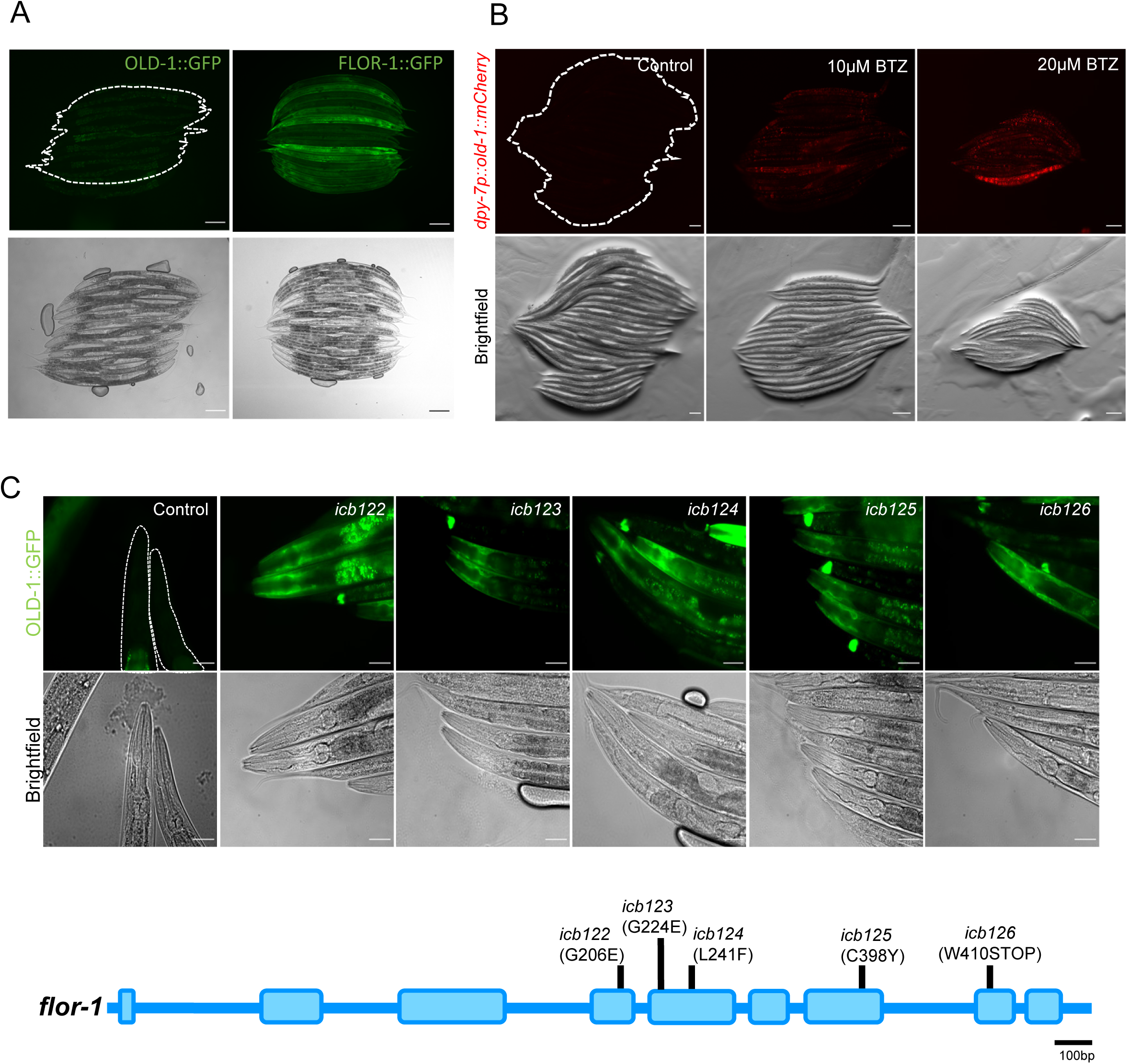
Inhibition of the proteasome and loss of function of *flor-1* causes OLD-1 accumulation. **(A)** Expression of OLD-1::GFP is low in comparison to FLOR-1::GFP. **(B)** Addition of the proteasome inhibitor Bortezomib to an *old-1::mCherry* expressing line (*icbEx356)* causes an increase of protein expression, suggesting OLD-1 is kept tightly regulated by the proteasome. Scale bar for A & B is 100 µm, n > 20. **(C)** Five additional *flor-1* alleles identified in an EMS suppressor screen all showed loss of *chil-27p::GFP* expression and an accumulation of OLD-1::GFP in the epidermis, indicating that FLOR-1 regulates OLD-1 levels at the epidermal membrane. Scale bar is 50 µm.

**Figure S5:**
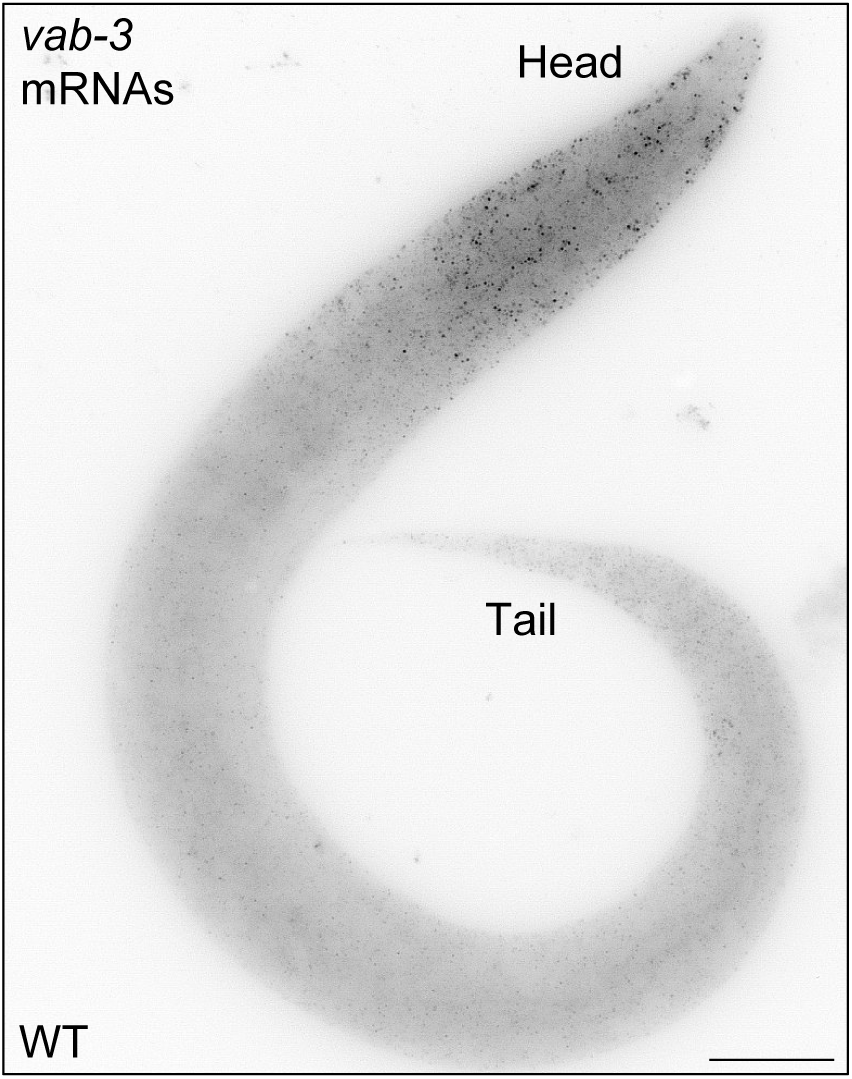
Expression pattern of *vab-3* in *C. elegans*. L1 stage animal showing anterior localization of *vab-3* mRNAs by smFISH. Scale bar is 10 µm.

## Supplementary Table Legends

**Table S1:** Differentially expressed genes upon extract treatment in N2, *old-1(-)* and *flor-1(-)* animals vs no extract treatments, and in animals overexpressing *old-1 (+ old-1p::old-1)* compared with non-transgenic animals.

**Table S2:** Differentially expressed genes upon epidermal overexpression of *old-1 (+ dpy-7p::old-1)* compared with WT animals carrying the co-injection marker (hygromycin resistance).

**Table S3:** Strains and oligos used in the study

